# Novel insights into *I*_Kur_ modulation by Lgi3-4: Implications in atrial fibrillation

**DOI:** 10.1101/2024.10.03.616587

**Authors:** Paula G. Socuéllamos, Álvaro Macías, Ángela de Benito-Bueno, Francisco M. Cruz, María Redondo-Moya, María José Coronado, Elvira Ramil, Silvia Rosado, Elsa Carolina Rios-Rosado, María Valencia-Avezuela, Laura de Andrés-Delgado, José Antonio Blázquez González, Alberto Forteza-Gil, Marta Gutiérrez-Rodríguez, José Jalife, Carmen Valenzuela

## Abstract

**Background:** Patients with atrial fibrillation (AF) exhibit a reduction in the ultrarapid outward potassium current (*I*_Kur_) conducted by K_V_1.5 channels. Ion channels are closely modulated by regulatory subunits, forming macromolecular complexes known as channelosomes. One such regulatory family is the leucine-rich glioma-inactivated protein family (Lgi1-4), which has been shown to interact with K_V_1, modifying their trafficking and/or biophysical properties in neurons. However, the expression and impact of these proteins in the heart is still unknown. We investigated the role of Lgi3-4 proteins in cardiac electrophysiology, focusing specifically on *I*_Kur_, and their potential contribution to the pathophysiology of AF.

**Methods:** We used three complementary biological systems, including heterologous COS-7, HEK297 and CHO cells, AAV-mediated cardiac-specific Lgi4 gene transfer in mice (Lgi4 mice), and human samples from patients in sinus rhythm and AF. Our multidisciplinary approach included immunolocalization, patch clamping, surface ECG, transvenous catheter-mediated intracardiac stimulation, and molecular biology techniques.

**Results:** Only Lgi3 and Lgi4 were expressed in the human heart. In human atrial tissue and heterologous cells, Lgi3 and Lgi4 interacted with K_V_1.5 channels. In HEK293 cells, Lgi3-4 impaired K_V_1.5/K_V_β association, partially reversing the K_V_β-induced N-type inactivation and reducing *I*_Kur_ amplitude. On surface ECG, the QRS interval was prolonged, and impulse conduction was impaired in cardiac-specific Lgi4 mice compared with control. In isolated ventricular cardiomyocytes from Lgi4 mice, early action potential repolarization was prolonged compared to control cardiomyocytes. These results correlated with the reduced K_V_1.5 membrane expression and *I*_Kur_ density observed in Lgi4 cardiomyocytes and HEK293 cells. Notably, Lgi4 protein expression was lower in atrial tissue from patients with AF than sinus rhythm patients. The reduction in Lgi4 protein levels in AF was also associated with an altered colocalization with K_V_1.5 channels, suggesting potential disruptions in their functional interactions.

**Conclusions:** Lgi3-4 proteins are new components of the K_V_1.5 channelosome. They modulate *I*_Kur_ by interfering with K_V_1.5 interaction with the K_V_β subunit. Importantly, Lgi4 is dysregulated differently in paroxysmal versus permanent AF. The results improved the understanding of this most common type of arrhythmia and identified Lgi proteins as a new potential target for treatment.

**NOVELTY AND SIGNIFICANCE:** What is known?

- Leucine-rich glioma-inactivated protein family (Lgi1-4) exert an important role in the nervous system and neurological diseases. In neurons, certain Lgi proteins interact with K_V_1 channels, modifying their trafficking and/or biophysical properties.
- In cardiomyocytes, the activation of K_V_1.5 channels generates the ultrarapid outward potassium current (*I*_Kur_), which is essential for the initial phase of human atrial repolarization, and it is dysregulated in AF.
- Changes in the properties or functional expression of some K_V_1.5 interacting proteins have crucial pathophysiological consequences.

What new information does this article contribute?

- We demonstrate that Lgi3-4 are novel components of K_V_1.5 channelosome, modulating *I*_Kur_ and hence human atrial electrophysiology. Lgi3-4 proteins decrease *I*_Kur_ by interfering with the interaction between K_V_1.5 and K_V_β subunits.
- The decrease in *I*_Kur_ in cardiac-specific mouse model expressing Lgi4 slows the early repolarization in the action potential, as well as produce electrophysiological changes in the surface ECG and the cardiac conduction system.
- Lgi4 is dysregulated differently in paroxysmal (PX) versus permanent (PM) AF, thus shedding light into the mechanisms underlying this cardiac arrhythmia.

## INTRODUCTION

K_V_1.5 channels, coded by *KCNA5,* generate the ultrarapid outward potassium current (*I*_Kur_), an atrial selective K^+^ current crucial for the initial phase of human atrial repolarization ^1–3^. *I*_Kur_ is altered in atrial fibrillation (AF), and is a potential target for the disease given its atrial selectivity. AF is the most common sustained arrhythmia and the most frequent cause of stroke ^2,4,5^. Both gain- and loss-of-function mutations in *KCNA5* have been associated with AF susceptibility ^6,7^. Furthermore, K_V_1.5 expression is reduced in the atria of AF patients compared with those in sinus rhythm (SR) ^3,8,9^. Anti-arrhythmic drugs are the first treatment option, but their effects are suboptimal, lacking sufficient efficacy and safety due to cardiac and extracardiac side effects ^2,10,11^. Even though several promising selective K_V_1.5 blockers have been developed in the last few years, no clinical trials have demonstrated a reduction of AF burden by K_V_1.5 inhibition. Therefore, whether *I*_Kur_ block is effective in the pharmacological AF conversion to SR, or reducing AF burden remains unanswered. Thus, new therapeutic approaches are urgently needed and may be developed, for example, by deciphering the molecular basis of *I*_Kur_ and its regulation in AF.

In human cardiomyocytes, K_V_1.5 channels assemble and form macromolecular complexes (channelosomes) with several K_V_β subunits (K_V_β1.2, K_V_β1.3 and K_V_β2.1) that induce: 1) fast and partial N-type *I*_Kur_ inactivation (only in the case of K_V_β1.x); 2) greater degree of C-type inactivation, 3) slower deactivation, and 4) negative shift of the activation curve ^12–14^. Also, K_V_β2.1 acts as a chaperone for K_V_1.5 in mouse ventricular myocytes, increasing *I*_Kur_ ^15^, but not in heterologous systems ^16^. However, the exact composition of these channelosomes remains undeciphered. Numerous interactors have been described recently, such as the membrane-associated Guanylate kinase (MAGUK) family (i.e. SAP97) ^17,18^, KChIP2 ^19^ or four-and-a-half Lim protein (FHL-1) ^20^. Changes in the myocardial ion channels or their interactors may produce dramatic alterations in action potential waveforms, synchronization, propagation and rhythmicity, thereby predisposing the heart to life-threatening arrhythmias ^2,21^.

The Lgi protein family includes four members (Lgi1-4) encoded by four genes (*LGI1-4*). These proteins consist of a leucine-rich repeat (LRR) domain in the N-terminal and an epilepsy-associated or epitempin (EPTP) domain in their C-terminal ^22,23^. Recent studies demonstrated that Lgis have complementary roles in different parts of the nervous system and cannot be replaced by another member of the Lgi family ^24,25^. Lgi1-4 proteins play critical regulatory functions by interacting with different members of the ADAM (A Disintegrin And Metalloproteinase) family, ADAM23, ADAM22 and ADAM11 ^26^. Lgi1 modulates synaptic transmission as the extracellular binding partner of ADAM22/23. Also, Lgi1 assembles with K_V_1.1, K_V_β1 and K_V_1.4 in neurons, selectively removing the N-type inactivation induced by the K_V_β1 subunit ^27^ and modulates the trafficking of K_V_1.1, K_V_1.2 and K_V_4.2 channels, tuning the action potential firing ^28–30^. Numerous mutations and deletions in the *LGI1* gene have been linked to autosomal dominant lateral temporal (lobe) epilepsy (ADLTE) ^22,28,30–33^. Additionally, antibodies against Lgi1 have been related to autoimmune encephalitis ^28,34–36^. Lgi3 modulates the K_V_1 channel complex localization at the juxtaparanodes (JXP) in myelinated axons in the central nervous system (CNS), this effect is mediated by ADAM23 ^24,37^. A cohort of patients carrying mutations in Lgi3 that display facial myokymia and developmental delay has been recently described ^37^. Lgi4 is an important regulator of myelination in the peripheral nervous system via its interaction with ADAM22, and its dysfunction results in peripheral hypomyelination ^25,38–41^. Mutations in Lgi4 have been linked to arthrogryposis multiplex congenita, with a very early mortality rate ^41,42^. However, the role of these proteins in the heart remains undeciphered.

Here, we explored the role of Lgi3-4 proteins in cardiac electrophysiology, specifically looking at their impact on *I*_Kur_ and how they might contribute to AF development. Using three different biological systems and a multidisciplinary approach, we have shown for the first time that Lgi3-4 proteins are new components of the K_V_1.5 channelosome. We discovered that they modulate *I*_Kur_ by interfering with the interaction between K_V_1.5 and the K_V_β subunits. Additionally, we found strong evidence that Lgi4 is dysregulated differently in paroxysmal versus permanent AF, shedding new light on the mechanisms underlying this type of cardiac arrhythmia.

## METHODS

### Data Availability

The data that supports the findings of this study are available from the corresponding author upon reasonable request.

### Ethics Statement

All human samples were obtained with the appropriate informed consent according to the World Medical Association Declaration of Helsinki principles of medical research involving human subjects and their use was approved by the Ethics Committee of the La Paz University Hospital (PI-2550). Approval was obtained also from the Ethics Committee of the Puerta de Hierro University Hospital (PI-158-22), the Ethics Committee in Human and Animal Experimentation CEEHA and the Biosecurity Committee of the CSIC (PI-2550 and 046/2023). All procedures were done under the 1527/2010 (November 15th, 2010) and the 1716/2011 (November 18th, 2011) Royal Decrees, as well as the 9/2014 (July 4th, 2014) Royal Decree-Law. All animal procedures conformed to Directive 2010/63/EU guidelines of the European Parliament on the protection of animals used for scientific purposes and to Recommendation 2007/526/EC, enforced in Spanish law under Real Decreto 53/2013. Animal protocols followed the Spanish National Center for Cardiovascular Research (CNIC) Institutional Ethics Committee recommendations and were approved by the Animal Experimentation Committee (Scientific Procedures) of Comunidad de Madrid (PROEX 111.4/20 and PROEX 226.5/23).

### Human samples

The Cardiac Surgery Service of the Health Research Institute of La Paz University Hospital (IdiPAZ) and Puerta de Hierro University Hospital provided human right atrial appendages. Ventricular tissue was kindly provided by the Departamento de Anatomía, Histología y Neurociencias of the Universidad Autónoma de Madrid (UAM).

### Cell culture

All cell lines were obtained from the American Type Culture Collection (Rockville, MD, USA) and were grown in their corresponding supplemented medium at 37°C in a 5% CO_2_ humidified atmosphere.

### Mice

C57BL/6J mice (5-6-weeks-old) were obtained from the Charles River Laboratories. Mice were reared and housed following CNIC institutional guidelines and regulations. The mice had free access to food and water.

### Adeno-Associated Virus Vector Production, Purification, and Mouse Model Generation

AAV vectors were generated encoding Lgi4 (NM_139284.3), followed by tdTomato reporter, or only tdTomato for the Control, under the expression of the cardiomyocyte-specific cTnT (cardiac troponin T) proximal promoter. Vectors were packaged into AAV serotype 9 and produced by the triple-transfection method, using HEK293T cells as described previously ^43,44^. Mice were anesthetized with ketamine (60 mg/kg) and xylazine (20 mg/kg) via the intraperitoneal route. Thereafter, 3.5×10^10^ virus particles were inoculated through the femoral vein in a final volume of 50 μL as previously described ^45^. All experiments were performed 8 to 10 weeks after infection.

### Cardiomyocyte Isolation

Mouse ventricular cardiomyocytes (CMs) were isolated as previously described ^46^. Ventricular CMs were used in this study because *I*_Kur_ and *I*_to_ constitute the main repolarizing currents in these cells.

### Patch Clamp

Whole-cell patch-clamp technique, internal and external solutions, ion currents, action potential acquisition, and data analysis were similar to those previously described for cell lines ^14,47^ and mouse ventricular cardiomyocytes ^46,48^.

### Protein extraction, Immunoprecipitation and Immunoblotting

HEK293 cells were homogenized with Lysis Buffer 1. Human atria tissue was mechanically homogenized with a Polytron (ULTRA-TURRAX® T10 Basic Disperser, IKA® Works) with Lysis Buffer 2. Protein content was determined by using a BCA Pierce Kit. We used Protein A or G Sepharose® beads incubated with the corresponding antibodies for coimmunoprecipitation assays. Protein extracts were separated by SDS-PAGE (8% acrylamide/bisacrylamide) gels, transferred to 0.45 μm PVDF membranes and incubated with different antibodies (See Supplementary material).

### Immunofluorescence

Different immunofluorescence protocols were used in COS-7 cells ^14^, cardiomyocytes ^46^ and human atria slices, as detailed in Supplementary material.

### Surface ECG Recordings

Mice were anesthetized using isoflurane inhalation (0.8%–1.0% volume in oxygen) and maintained at 37 °C on a heating plate. Four-lead surface ECGs were recorded for 5 minutes using subcutaneous limb electrodes connected to an MP36R amplifier(BIOPAC Systems). Data acquisition and analysis were performed using the AcqKnowledge software.

### In Vivo Intracardiac Recording and Stimulation

An octopolar catheter (Science) was inserted through the jugular vein and advanced into the right atrium and right ventricle as previously described ^49^. Atrial and ventricular arrhythmia inducibility was assessed by applying consecutive S1 and S2 pulse trains at 10 and 25 Hz, respectively.

### Statistical Analyses

Data are expressed as mean ± SEM of n experiments, where N represents the number of patients or animals, and n represents the number of individual cells, IF images or cell lysates. Comparisons were performed between different experimental groups by an unpaired two-tailed Student’s t-test or using multiple comparison Mann-Whitney’s test. When more than two experimental groups were compared, one-way or two-way ANOVA with Tukey’s or Šídák’s multiple comparison test, respectively, were used. Contingency analysis was performed with chi-square test. Differences were considered significant when p<0.05.

## RESULTS

### Lgi protein expression and co-localization with K_V_1.5 channels in the human heart

To determine the expression pattern of Lgi proteins in human myocardium, we performed immunofluorescence analysis in samples from the right atrium and right ventricle. Only Lgi3 and Lgi4 were expressed in human atrial and ventricular cardiomyocytes, whereas Lgi1 and Lgi2 were nearly absent (Figure 1A). Double staining of Lgi3-4 and K_V_1.5 revealed the colocalization of Lgi3 (upper panels) and Lgi4 (lower panels) with K_V_1.5 in the human atrium (Figure 1B) and COS-7 cells (Figure 1C) with a Pearson’s correlation coefficient (PCC) of about 0.5 and 0.8, respectively. Lgi3-4 were expressed mainly in the endoplasmic reticulum (ER) and plasma membrane (Figure 1C, Figure S1), as previously reported ^38^. Coimmunoprecipitation studies confirmed the interaction of Lgi3-4 with K_V_1.5 (Figure 1D).

**Figure 1.**
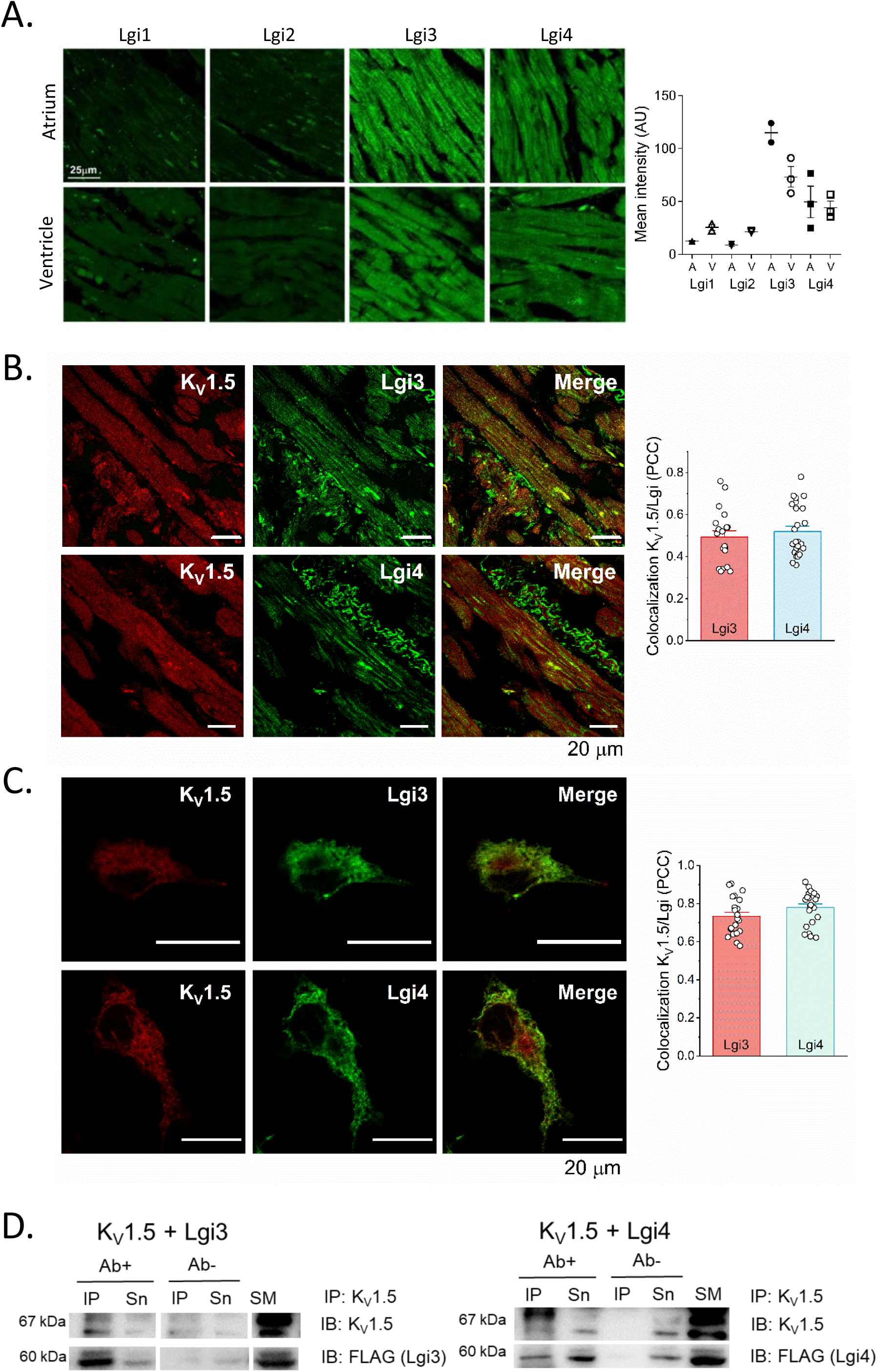
Lgi3-4 are expressed in human myocardium and interact with K_V_1.5. **A)** Confocal images of the immunodetection of Lgi1-4 in human atrium and ventricle slices obtained from healthy patients. Right graph: quantification of the Lgis expression in atrium (A) and ventricle (V). The intensity mean was measured as the integrated density. Values represent mean±SEM of N=2-3 donors, n=9. **B)** Representative confocal images of the immunodetection of K_V_1.5 along with Lgi3 (upper panels) and Lgi4 (lower panels) in human atrium from patients in SR (N=4-5, n=20-25). The Pearson’s correlation coefficient (PCC) between either Lgi3-4 and K_V_1.5 is shown at the right graph. Values represent mean±SEM. **C)** Representative confocal images of COS7 cells cotransfected with K_V_1.5 and Lgi3 (upper panels) or Lgi4 (lower panels), with the PCC shown in the right bar graph. Data represent mean±SEM of n=22-23. **D)** Coimmunoprecipitation of K_V_1.5 with Lgi3 (left) or Lgi4 (right) coexpressed in HEK293 cells, using the anti-K_V_1.5 antibody (IP: K_V_1.5) and immunoblotting (IB) K_V_1.5 and FLAG (Lgi3-4) (n=3). SM: Starting Material, IP: Immunoprecipitated, Sn: Supernatant, IB: Immunoblotting. In all experimental conditions, at least 3 independent experiments were performed.

### Lgi3-4 decrease the amplitude of K_V_1.5/K_V_β current, but not K_V_1.5 alone

To investigate the functional impact of Lgi3-4 proteins on K_V_1.5 electrophysiology, we conducted experiments in HEK293 cells expressing Lgi3-4 and co-transfected with K_V_1.5 alone or in the presence of different cardiac K_V_β subunits (K_V_β2.1, K_V_β1.3 and K_V_β1.2). Lgi3 and Lgi4 had no significant effect on the K_V_1.5 current amplitude (Figure 2A and 2B). However, when K_V_1.5 channels were assembled with K_V_β2.1, K_V_β1.2 or K_V_β1.3 subunits, Lgi3-4 exerted two striking effects on K_V_1.5 currents: 1) the effects of K_V_β2.1 on C-type inactivation and those of K_V_β1 on N-type inactivation were almost abolished in the presence of Lgi3-4 (Table S1, Figure 2C, 2E and 2G), resembling the current elicited by K_V_1.5 channels alone; and 2) the current amplitude of K_V_1.5 coexpressed with K_V_β2.1, K_V_β1.3 and K_V_β1.2 was reduced when Lgi3-4 were expressed, both at the end of the depolarizing pulse and at the peak amplitude (Figure 2D, 2F and 2H). The effects on current amplitude could be due to changes in the trafficking of K_V_1.5 channels. Flow cytometry experiments showed that Lgi3-4 decrease the membrane expression of K_V_1.5 in the presence of K_V_β2.1, K_V_β1.2 or K_V_β1.3 (Figure 2I), but not in the absence of K_V_β subunits. These results could explain, at least in part, the reduction observed in the current amplitude. Another explanation for this could be a shift in the voltage dependence of activation towards more positive potentials. Lgi4, but not Lgi3, shifted K_V_1.5 V_h_ of activation towards more negative potentials. For K_V_1.5/K_V_β2.1 currents, either Lgi3-4 shifted the V_h_ towards more positive potentials, which is in agreement with a reduction in the current amplitude. Neither Lgi3 nor Lgi4 changed the V_h_ of K_V_1.5/K_V_β1.3 and K_V_1.5/K_V_β1.2 currents (Table S2).

**Figure 2.**
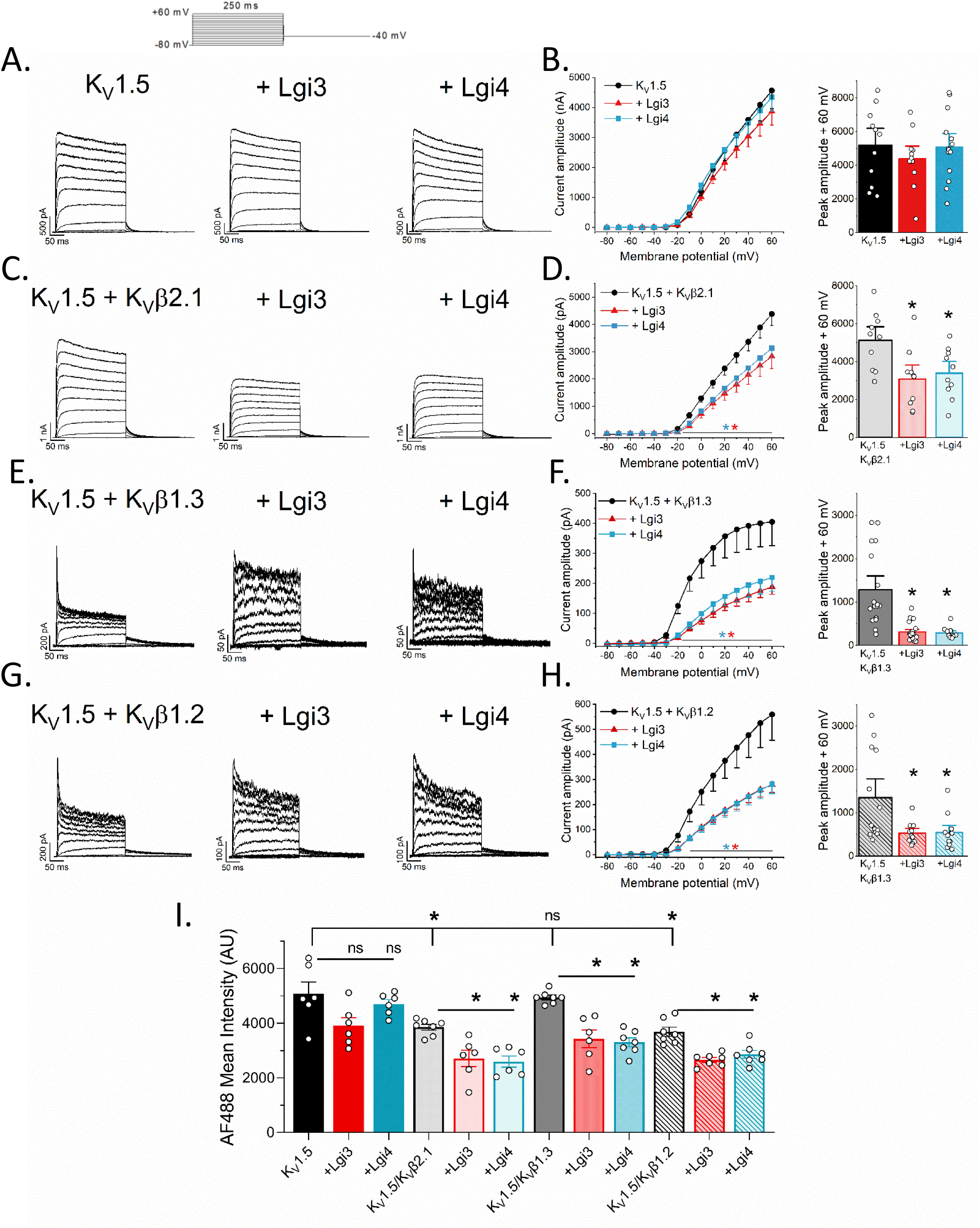
Electrophysiological effects of Lgi3-4 on K_V_1.5 currents. ***Left panels***: Representative current traces elicited after the application of the I-V protocol shown in the upper panel in HEK293 cells transfected with K_V_1.5 with or without Lgi3 or Lgi4 (**A**) and in the presence of K_V_β2.1 (**C**), K_V_β1.3 (**E**) or K_V_β1.2 (**G**) subunits. Note that K_V_1.5/K_V_β currents are more similar to K_V_1.5 currents alone when Lgi3 or Lgi4 are expressed. ***Right panels***: I-V relationships measured at the end of the 250 ms depolarizing-pulse and current amplitude at +60 mV measured at the peak in K_V_1.5 with or without Lgi3 (red) or Lgi4 (blue) (n=11-14) (**B**), and in the presence of: K_V_β2.1 (n=10) (**D**), K_V_β1.3 (n=14-24) (**F**) or K_V_β1.2 (n=10-13) (**H**) subunits. (**I**) Membrane expression of K_V_1.5 in HEK293 cells transfected with K_V_1.5-HA with or without Lgi3 (red) or Lgi4 (blue) and in the presence of K_V_β2.1, K_V_β1.3 or K_V_β1.2 subunits (n=6-7). Values represent mean±SEM of the indicated n. (Non-paired t-test \**p* < 0.05).

### Lgi3-4 reduce K_V_β-induced inactivation through competitive binding

As stated above, Lgi3-4 reduce the effects induced by K_V_β subunits on the inactivation of K_V_1.5 channels. To better understand why, we used various electrophysiological protocols, taking into account that the N-type inactivation is highly voltage-dependent ^14,50^. First, we applied depolarizing pulses at +60 and at +140 mV (Figure 3A). The traces elicited at +140 mV enabled us to analyze the kinetics of inactivation obtaining the time constants (τ_fast_ and τ_slow_, for the fast or N-type and slow or C-type inactivation, respectively), and the relative contribution of the slow component of inactivation (A_slow_), which corresponds to the A_slow_/(A_slow_ and A_fast_).

**Figure 3.**
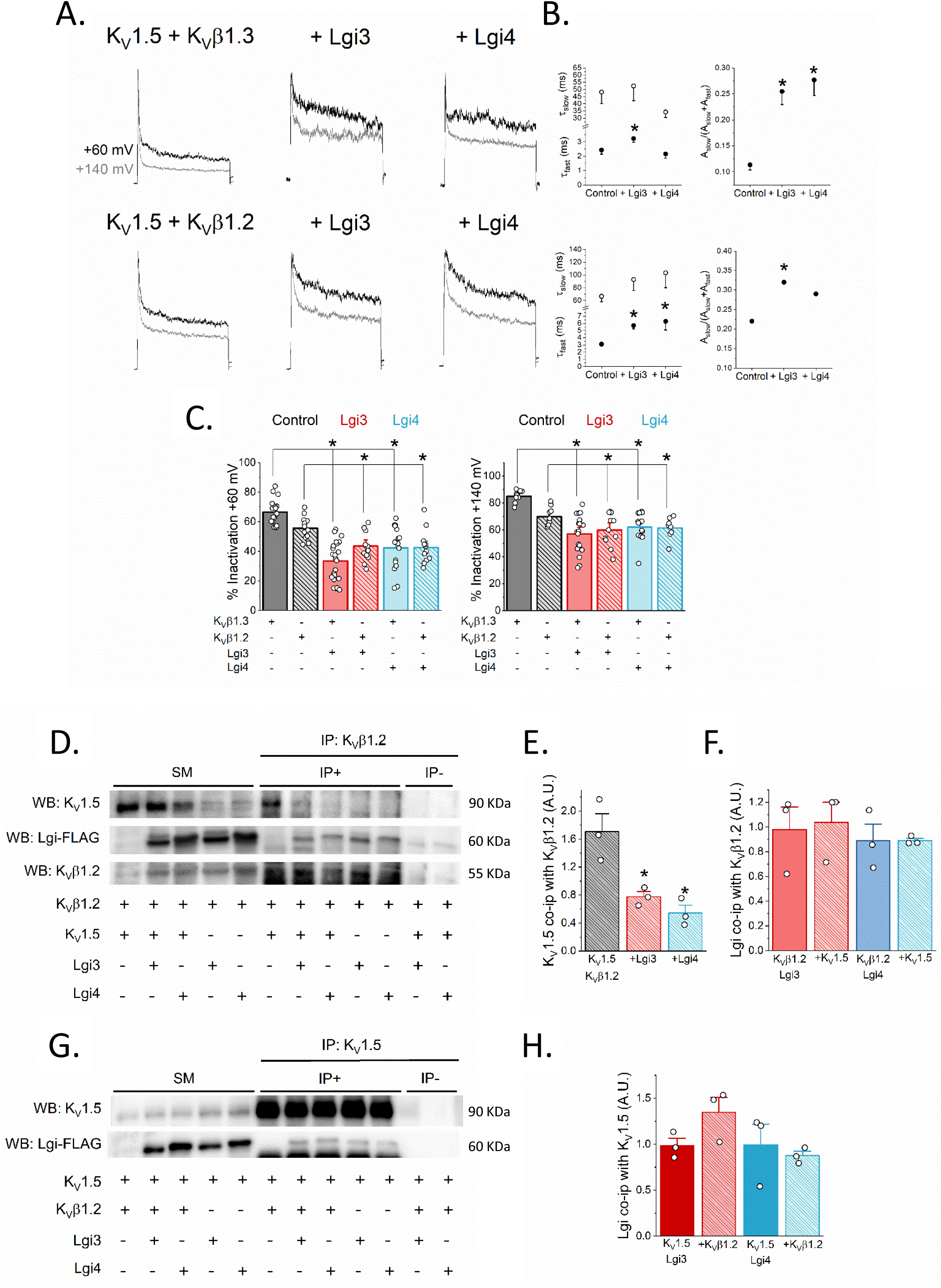
Lgi3-4 decrease the N-type inactivation induced by K_V_β1 on K_V_1.5 currents and hinder the association between K_V_1.5 and K_V_β. **A**) Representative current traces recorded after application of 250 ms depolarizing pulses to +60 (black) or +140 mV (grey) in HEK293 cells transfected with K_V_1.5/K_V_β1.3 (upper panel) or K_V_1.5/K_V_β1.2 (lower panel) in the absence and in the presence of Lgi3 or Lgi4. **B**) Inactivation kinetics measured at +140 mV fitted to a double exponential. The left graphs show the time constant of inactivation and the right graphs show the contribution of the slow inactivation to the whole inactivation process (n=8-18). **C**) Degree of inactivation of K_V_1.5 with (+) or without (-) K_V_β1.3, K_V_β1.2, Lgi3 and/or Lgi4 currents at +60 mV (left graph, n=12-23) and +140 mV (right graph, n=8-18). Values represent mean±SEM. Non-paired t-test were performed comparing each condition with its corresponding control. \**p* < 0.05. **D**) HEK293 cells were cotransfected with (+) or without (-) K_V_1.5, K_V_β1.2, Lgi3 and Lgi4. Total lysates were immunoprecipitated (IP) against K_V_β1.2 and immunoblotted (IB) against K_V_1.5, FLAG (Lgi3-4) and K_V_β1.2 (n=3). **E**) Quantification of K_V_1.5 coimmunoprecipitation with K_V_β1.2 measured as K_V_1.5 expression in the IP+ divided by its corresponding SM expression. Note that K_V_1.5 colocalization with K_V_β1.2 decreases when Lgi3-4 are coexpressed. **F**) Quantification of Lgi3-4 (FLAG) coimmunoprecipitation with K_V_β1.2 measured as FLAG expression in the IP+ divided by its corresponding SM expression. **G**) Total lysates were immunoprecipitated (IP) against K_V_1.5 and immunoblotted (IB) versus K_V_1.5 and FLAG (Lgi3-4) (n=3). (**H**) Quantification of FLAG coimmunoprecipitation with K_V_1.5 measured as FLAG expression in the IP+ divided by its corresponding SM expression. Values represent mean±SEM of the n indicated above. Non-paired t-test were performed comparing each condition with its corresponding control (\**p* < 0.05). SM: starting material, IP+: IP in the presence of antibody, IP-: IP in the absence of antibody.

We tested whether Lgi3 and Lgi4 decrease the contribution of the N-type inactivation either by: **1)** slowing τ_fast_ or **2)** increasing the A_slow_ contribution. Neither Lgi3 nor Lgi4 modified the τ_slow_ values in any experimental condition (Figure 3B). Lgi3-4 clearly reduced the degree of inactivation of K_V_1.5/K_V_β1.3 and K_V_1.5/K_V_β1.2 currents at either membrane potential (Figure 3C). These effects can be explained if: **1)** Lgi3-4 shift the voltage dependence of N-type inactivation to more positive potentials, or **2)** Lgi3-4 hinder the interaction of K_V_1.5 with K_v_β1. To test the first hypothesis, we analyzed the effects of Lgi3-4 on the voltage-dependence of N-type inactivation induced by K_V_β1.3 or K_V_β1.2, which was only slightly modified (Table S3, Figure S2). The degree of inactivation accumulated during the 10-ms prepulse decreased strongly in the presence of Lgi3 or Lgi4, which was consistent with the above-mentioned reduced degree of N-type inactivation. Next, we conducted co-immunoprecipitation experiments to explore whether the effects of Lgi3-4 on K_V_1.5/K_V_β1 currents were due to competition between these distinct proteins. As shown in Figure 3D-E, K_V_1.5 coimmunoprecipitation with K_V_β1.2 was dramatically reduced in the presence of Lg3-4. Additionally, we found that Lgi3-4 bind to K_V_β1.2 and such interaction is disaffected by the presence of K_V_1.5 (Figure 3D and 3F). Finally, the presence of K_V_β1.2 did not modify the degree of co-immunoprecipitation between K_V_1.5 and Lgi3-4 FLAG-tagged (Figure 3G-H). These above results suggest that Lgi3-4 compete with K_V_β1.2 for its binding to K_V_1.5 channels; and also, that Lgi3-4 bind to both K_V_1.5 and K_V_β1.2. Altogether, the results suggest more complex mechanisms in the regulation of the macromolecular K_V_1.5 channelosome, extending beyond the scope of this study. The effects of Lgi3-4 on K_V_1.5/K_V_β1.3 currents were similar when the experiments were performed in cells transfected with ADAM23 (Figure S3).

### Cardiac-specific Lgi4 mice have abnormal ECGs and cardiac conduction

To study the biological contribution of Lgi proteins in vivo, we first analyzed the expression pattern of K_V_1.5, Lgi3 and Lgi4 in ventricular cardiomyocytes (CMs) (Figure S4A). Lgi3, but not Lgi4, was detected and widely distributed, and K_V_1.5 was expressed in mouse ventricular CMs in agreement with previous reports ^51,52^. Therefore, we used adeno-associated virus (AAV) technology ^45^ to generate a mouse model with cardiac-specific expression of Lgi4 under cTnT promoter, and demonstrated Lgi4 expression in Lgi4-transduced CMs (Figure 4A).

**Figure 4.**
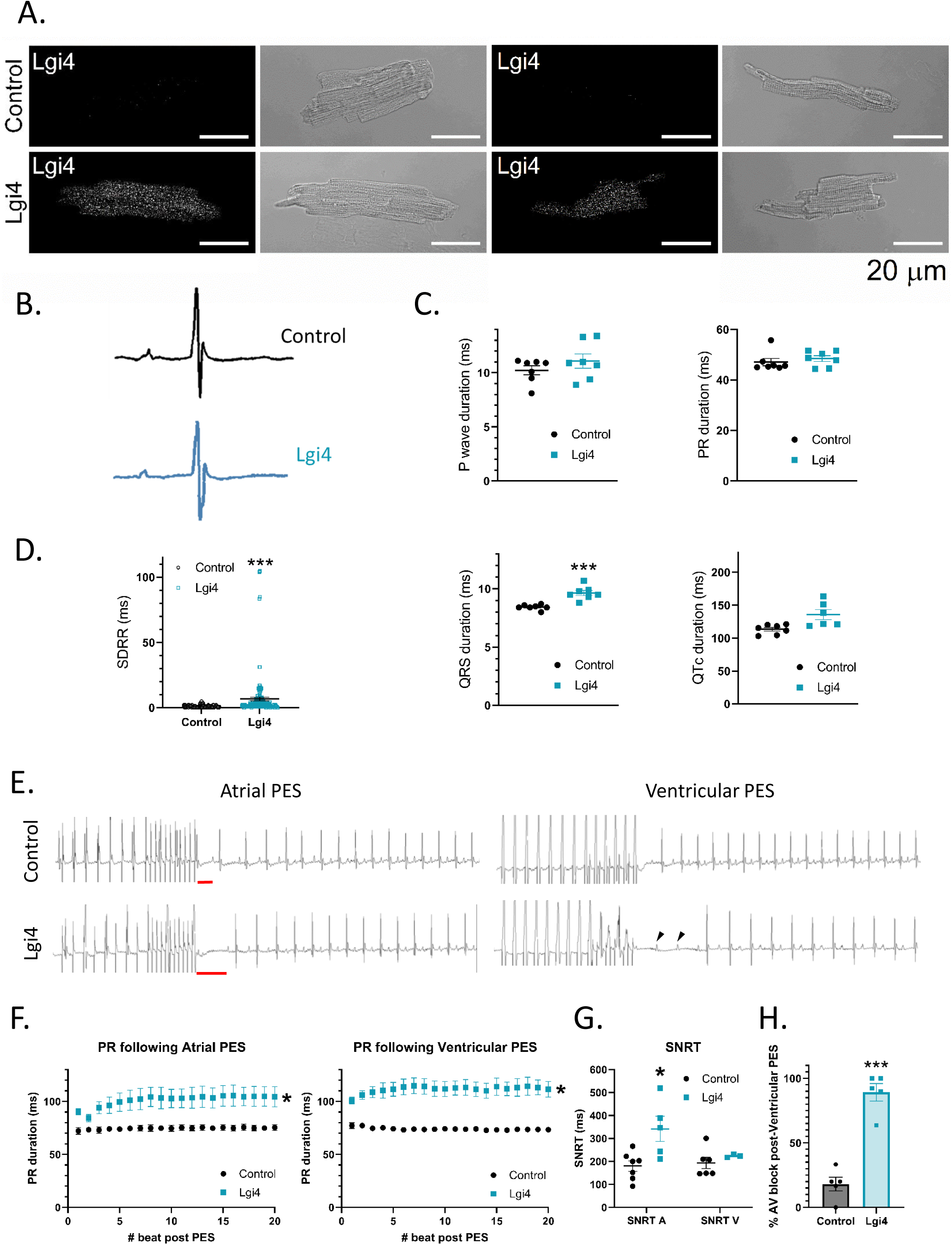
Lgi4 mice have abnormal ECGs and cardiac conduction. **A**) Representative examples of the immunodetection of Lgi4 in permeabilized mice ventricular CMs infected with Control (upper panels) or with Lgi4 expressing (lower panels) AAV9 (N=3, n=30). The corresponding bright field images are shown next to each immunostaining. **B**) Representative ECG traces of Control (black) and Lgi4 (blue) mice. **C**) P wave, PR, QRS and QTc intervals in control (black) and Lgi4 (blue) mice. Note that the QRS is prolonged in ECG from Lgi4 mice. **D**) Beat to beat variability is represented as the standard deviation of the RR (SDRR). Data are represented as mean±SEM of N=7 (Non-paired t-test \*\*\**p* < 0.001). **E)** Original lead-II ECG recordings during and after atrial (A) and ventricular (V) PES in Control (above) and Lgi4 (below) mice. The SNRT is shown in red and AV blocks are indicated with arrows. **F**) PR duration after atrial (left) and ventricular (right) stimulation in Control (black) and Lgi4 (blue) mice. **G**) Sinus node recovery time (SNRT) after atrial (A) and ventricular (V) stimulation in Control (black) and Lgi4 (blue) mice. Each point represents the mean of approximately 10 PES in each mice. **H**) Percentage of atrial-ventricular block following ventricular PES in Control (Black) and Lgi4 (blue) mice. Data are represented as mean±SEM (N=5-7). Non-paired t-test \**p* < 0.05, \*\*\**p* < 0.001.

We first studied the effects of cardiac-specific Lgi4 expression on surface-ECG lead-II parameters (Figure 4B). Lgi4 expression did not modify the P wave, PR and QTc intervals. However, the QRS interval was significantly longer in Lgi4 than control mice (Figure 4C). On the other hand, RR variability was increased in the Lgi4 model, as shown by the increased RR standard deviation (SDRR) (Figure 4D).

To determine whether cardiac Lgi4 expression is arrhythmogenic in anesthetized mice we used transvenous intracardiac programmed electrical stimulation (PES) in Lgi4 and control mice (Figure 4E). No atrial or ventricular arrythmias were detected in either group. However the PR interval substantially prolonged after atrial and ventricular PES in Lgi4 compared with control mice (Figure 4F), which may indicate that atrioventricular (AV) node or Purkinje fiber conduction was altered. In addition, the sinus node recovery time (SNRT) was prolonged in Lgi4 mice when stimulating the atria, but not the ventricle, in comparison to control mice (Figure 4G), which may indicate alterations in the sinoatrial node (SAN). Additionally, in contrast to control, ventricular PES led to long-lasting PR prolongation in Lgi4 mice exhibited AV block after most ventricular PES trains (Figure 4H), again suggesting Lgi4 effects on Purkinje fibers. Altogether, these results together with the substantially increased RR variability strongly indicate that cardiac expression of Lgi4 is detrimental to the cardiac conduction system.

### Lgi4 modulates APs properties in mouse cardiomyocytes

To determine whether Lgi4 affects action potential (AP) parameters like depolarization or early repolarization ^53^, we next set out to study the effects of Lgi4 expression on the action potential (AP) characteristics. Figure 5A shows original cardiac ventricular AP recorded from control and Lgi4 mice. Lgi4 expression did not modify the resting membrane potential (RMP), amplitude and maximal upstroke velocity (dV/dt_max_) (Figure 5B). Conversely, APD (measured at 20% to 70% repolarization) was longer in Lgi4 than control CMs, compatible with a slower early repolarization. However, APD_90_, which accounts for the total duration of repolarization, was not modified (Figure 5C). Changes in the AP waveform in Lgi4-transduced CMs can have different functional consequences ^54^. As illustrated by the representative recordings in Figure 5D, unlike control (top), during the application of a train of pulses at 5 Hz Lgi4 CMs showed progressive APD prolongation, as quantitated by the recovery curve (left) and the Poincaré plot (right) of Figure 5E. In addition, as shown by the lower panel of Figure 5D,F, at a certain point during the 5-Hz train some Lgi4 CMs exhibited excessive APD prolongation and early afterdepolarizations (EADs) that interfered with the stimulation. Figure 5F-G shows the quantification of these changes in control and Lgi4 CMs. In Figure 5F, the percentage of control and Lgi4 CMs exhibiting excessive APD prolongation plus EADs increases substantially when stimulating at 10 than 5 Hz. In Figure 5G it is shown that the first pulse with such features happened earlier in Lgi4 than control CMs.

**Figure 5.**
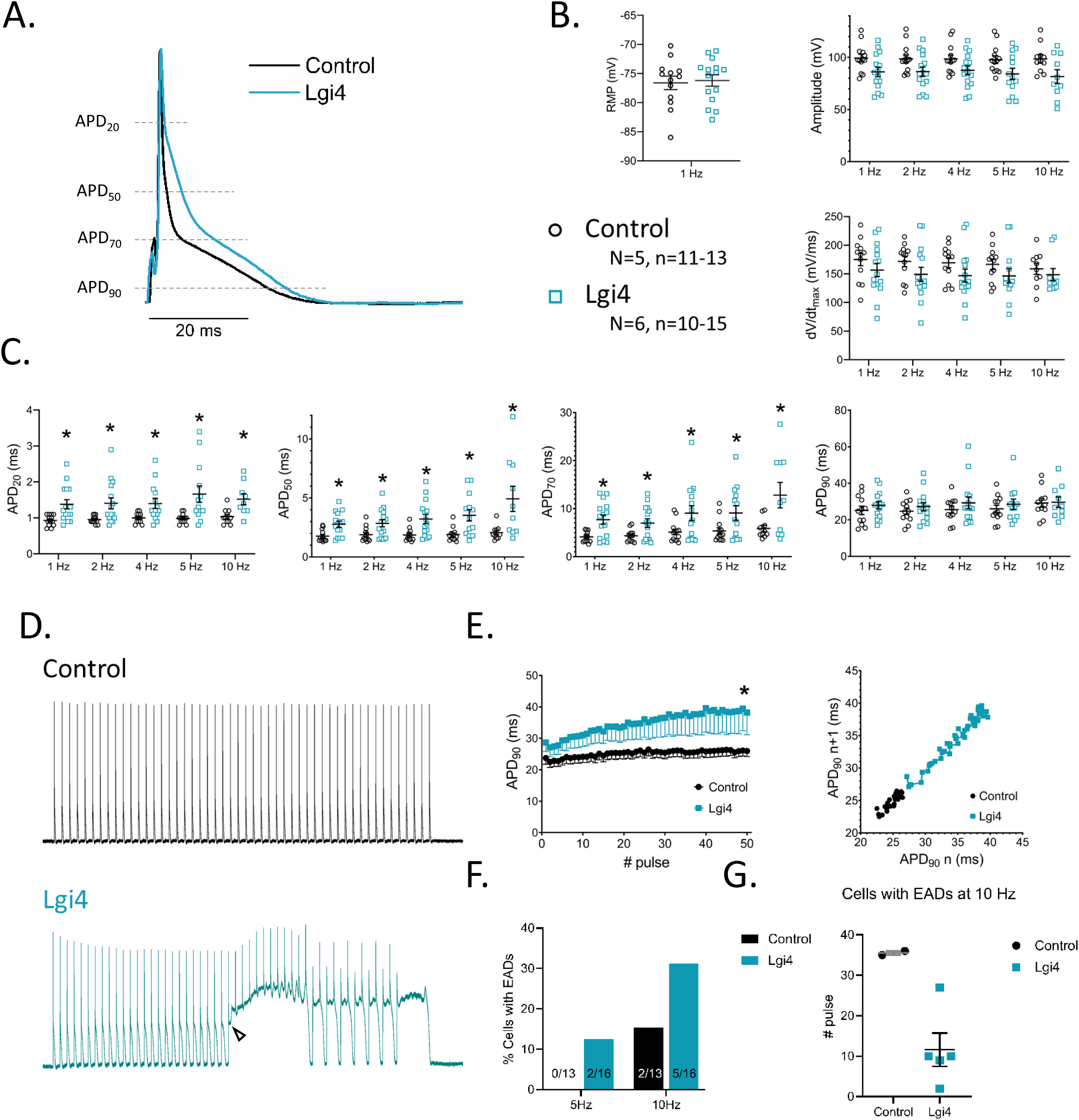
Lgi4 modulates early repolarization and induces APD prolongation and EADs at high stimulation frequencies. **A**) Representative recordings of APs registered at 1 Hz of stimulation in Control (black) and Lgi4 (blue) mice ventricular CMs. **B**) In the upper right graph, the resting membrane potential (RMP) at 1 Hz, the maximal amplitude of the AP (Amplitude) and the maximum upstroke velocity (dV/dt_max_) are represented; these two latter at different frequencies of stimulation. **C**) APD measured at the 20, 50, 70 and 90% of repolarization. Note that APD_20_ to APD_70_ are longer in CMs from Lgi4 mice than in Control ones. **D)** Original recordings of APs generated when stimulated at 5 Hz in Control (black) and Lgi4 (blue) ventricular CMs. Note the continuous depolarization in Lgi4 CMs. **E)** APD_90_ variability in Control (black) and Lgi4 (blue) when the CMs are stimulated at 5 Hz. **F)** Percentage of cells exhibiting progressive AP prolongation and EADs when stimulated at 5 and 10 Hz, and **G)** Number of the pulse in which the first EADs appear at 10 Hz, respectively, in control (black) and Lgi4 (blue) CMs. Values are represented as mean±SEM (N=5-6, n=10-15). (Two-tailed t-tests (A-C), mixed effects analysis and chi-square test (E-F). \**p* < 0.05).

### Lgi4 reduces *I*_Kur_ and K_V_1.5 membrane expression in ventricular CMs

We hypothesized that the reduction in early repolarization in CMs is due to a decrease in *I*_Kur_ previously demonstrated in heterologous systems (Figure 2). Considering that K_V_β2.1 and K_V_β1.2 are present in mouse CMs, we measured the repolarizing K^+^ currents in our AAV-mediated mouse models (Figure S5). Voltage-clamp experiments showed a significant *I*_Kur_ reduction in Lgi4 compared to control CMs (Figure 6A). *I*_to_ amplitude showed tendency to increase in Lgi4-CMs (Figure 6B). Because *I*_to_ is generated by the activation of K_V_4.3/K_V_4.2, its tendency to increase in our experiments might be due to differential Lgi4 modulation on each K_V_4 channel subtype. We tested this hypothesis in CHO-K1 cells transfected with K_V_4.3/Lgi3-4 or K_V_4.2/Lgi3-4 (Figure S6A-D). The experiments revealed that Lgi3-4 increased the current amplitude of K_V_4.3, but not K_V_4.2 channels, without changing their voltage dependence. In additional experiments, both Lgi3 and Lgi4 colocalized with K_V_4.3 with a PCC of about 0.4 in human atrium (Figure S6E) and with a PCC of 0.6 in COS-7 cells (Figure S6F). *I*_ss_ (generated by the activation of K_V_2.1) and the inward current (mainly driven by the activation of K_ir_2.1) were not modified in Lgi4-CMs versus control cells (Figure 6C-D). Immunofluorescence assays in non-permeabilized CMs demonstrated reduced expression of K_V_1.5 in the plasmalemma in Lgi4-expressing CMs (Figure 6E), which explains the reduction of *I*_Kur_. While Lgi4 did not modify total K_V_1.5 expression, it shifted its cellular distribution from the plasmalemma to intracellular compartments (Figure S7A). Next, we studied whether Lgi4 modified K_V_4.3 expression or cellular distribution, which did not occur (Figure S8A), even though these proteins colocalize with K_V_4.3 (Figures S8B-C). To sum up, Lgi4 reduces *I*_Kur_, likely due to a reduction in K_V_1.5 membrane expression.

**Figure 6.**
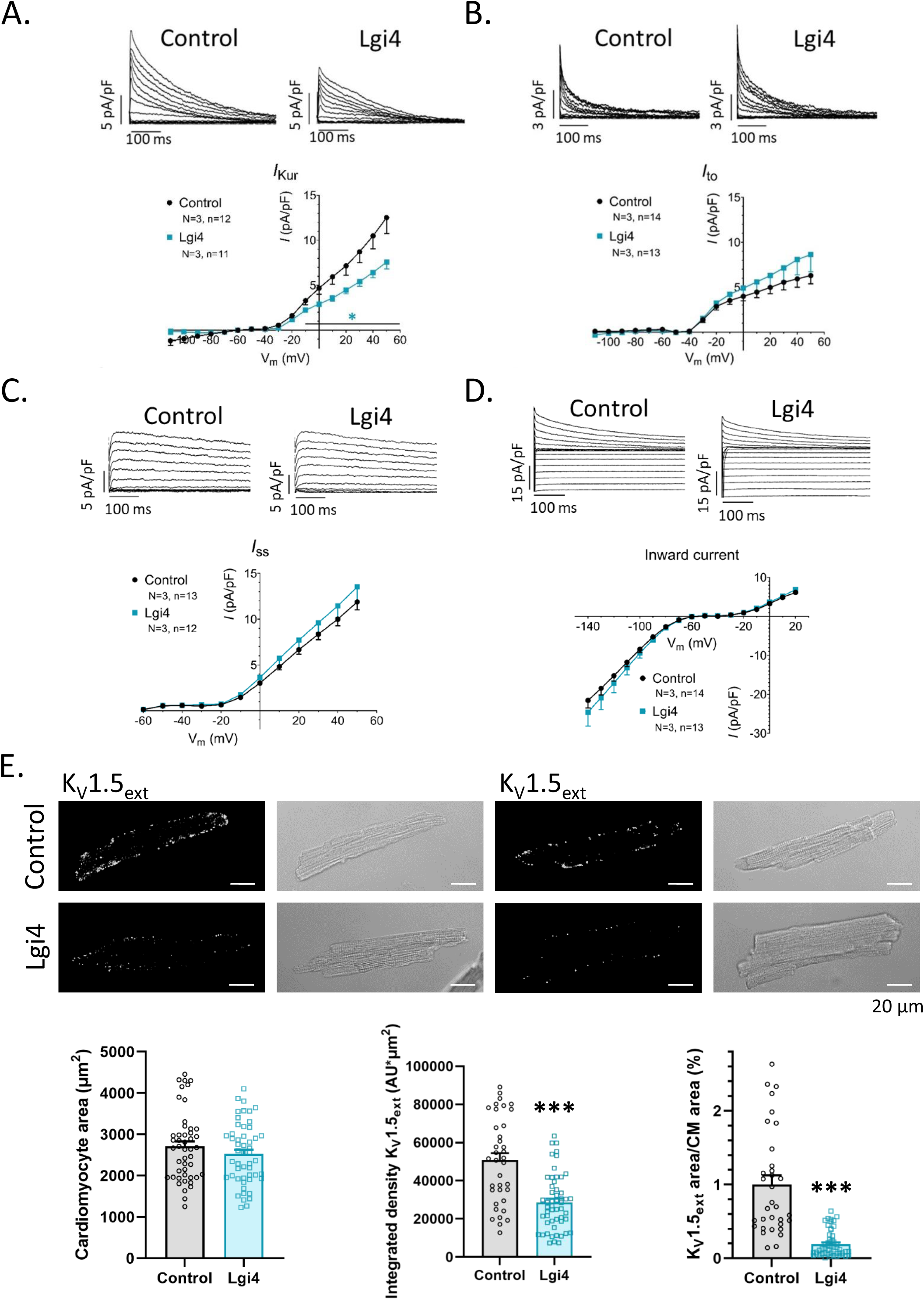
Lgi4 decreases *I*_Kur_ current amplitude and K_V_1.5 plasmalemmal expression in mice ventricular CMs. Original current traces of the outward potassium currents *I*_Kur_ (**A**), *I*_to_ (**B**) and *I*_ss_ (**C**) and the inward K^+^ current (**D**) recorded in Control (left) and Lgi4 (right) mice ventricular CMs, together with the current-amplitude (I-V) relationship for each current measured at the maximum peak amplitude in *I*_Kur_ (**A**), *I*_to_ (**B**), and *I*_ss_ (**C**) or at the end of the pulse in the inward K^+^ current (**D**) in mice ventricular CMs Control (black) and Lgi4 (blue). Data are represented as mean±SEM (N=3, n=11-14). Non-paired t-test were performed comparing each condition with its corresponding control. \**p* < 0.05. (**E**) Representative confocal images of the immunodetection of extracellular K_V_1.5 in non-permeabilized mice ventricular CMs Control and Lgi4. The corresponding bright field images are shown next to each immunostaining. Images clearly show that K_V_1.5 is less expressed in the plasmalemma in Lgi4 CMs than in Control ones. The quantification of the CMs area (left), the integrated density of K_V_1.5_ext_. Both analyses show that K_V_1.5 membrane expression is decreased in Lgi4 CMs. Data are represented as mean± SEM (N=4, n=33-53). Non-paired t-test \*\*\**p* < 0.001.

Adrenergic stimulation strongly modulates of human *I*_Kur_. Therefore, we compared the effects of isoprenaline on APs and *I*_Kur_ of Lgi4 versus control CMs. Figure S9A shows representative AP traces at 2 Hz before and after perfusion with isoprenaline 1 μM for 5-10 min (shown in discontinuous lines), as previously described ^55^. Isoprenaline did not change the RMP, maximal amplitude or upstroke velocity (upper panels). Conversely, the early repolarization of the APs (APD_50_ to APD_70_) became longer in the presence of isoprenaline in control than in Lgi4 CMs (lower panels). Slower repolarization could be due to an increase in *I*_CaL_ or a decrease in *I*_Kur_. Isoprenaline increases both current amplitudes in human atrial myocytes ^55,56^. Isoprenaline did not change the *I*_Kur_ amplitude in control CMs, whereas it augmented it in Lgi4 expressing CMs (Figure S9C). As Lgi4 CMs exhibit a smaller *I*_Kur_, its increase is greater, maybe because isoprenaline induce the plasmalemmal expression of K_V_1.5. These changes in *I*_Kur_ amplitude could explain why the APs after perfusion of control and Lgi4 CMs with isoprenaline are very similar, as well as the occurrence of continuous depolarizations (Figure S9B-C).

### K_V_1.5 channelosome is altered in atrial fibrillation

Human samples from the right atrial appendage were collected from patients under SR (control) and with AF in different stages of its progress. Table S4 shows the information of the patients included in this study. *KCNA5* mRNA levels are decreased in AF (Figure S10A) mostly due to a reduction in paroxysmal (PX) AF when compared to SR (Figure 7A). *LGI3* mRNA levels are decreased in PX and permanent (PM) AF, whereas only a tendency towards its reduction was revealed in *LGI4* mRNA levels in PM AF samples (Figure 7A). A positive correlation between *LGI3*/*KCNA5*, *LGI4*/*KCNA5* and *LGI3*/*LGI4* mRNA expression were observed (Figure S10D). Either the total protein expression of K_V_1.5, Lgi3 and Lgi4 in atrial homogenates from patients in SR or AF (Figure 7B, S10B) or the expression of these proteins in cardiomyocytes (Figures 7C *left*, 7D, S10C) followed the same trend. K_V_1.5 and Lgi4 protein expression was reduced in atrial homogenates from patients in PM AF, but not in PX AF; whereas Lgi3 protein expression was not modified in any case (Figure 7B). Both Lgi3 and Lgi4 protein expression exhibited a correlation trend with K_V_1.5 protein expression (Figure S10E). Unexpectedly, Lgi3 and Lgi4 had a positive correlation (Figure S10E). Surprisingly, the interaction between K_V_1.5 and Lgi4 was reduced in PX AF in comparison to SR, but not in PM AF, which can contribute to fine-tune the mechanisms involved in the AF electrical remodeling (Figure 7C *right*, 7E).

**Figure 7.**
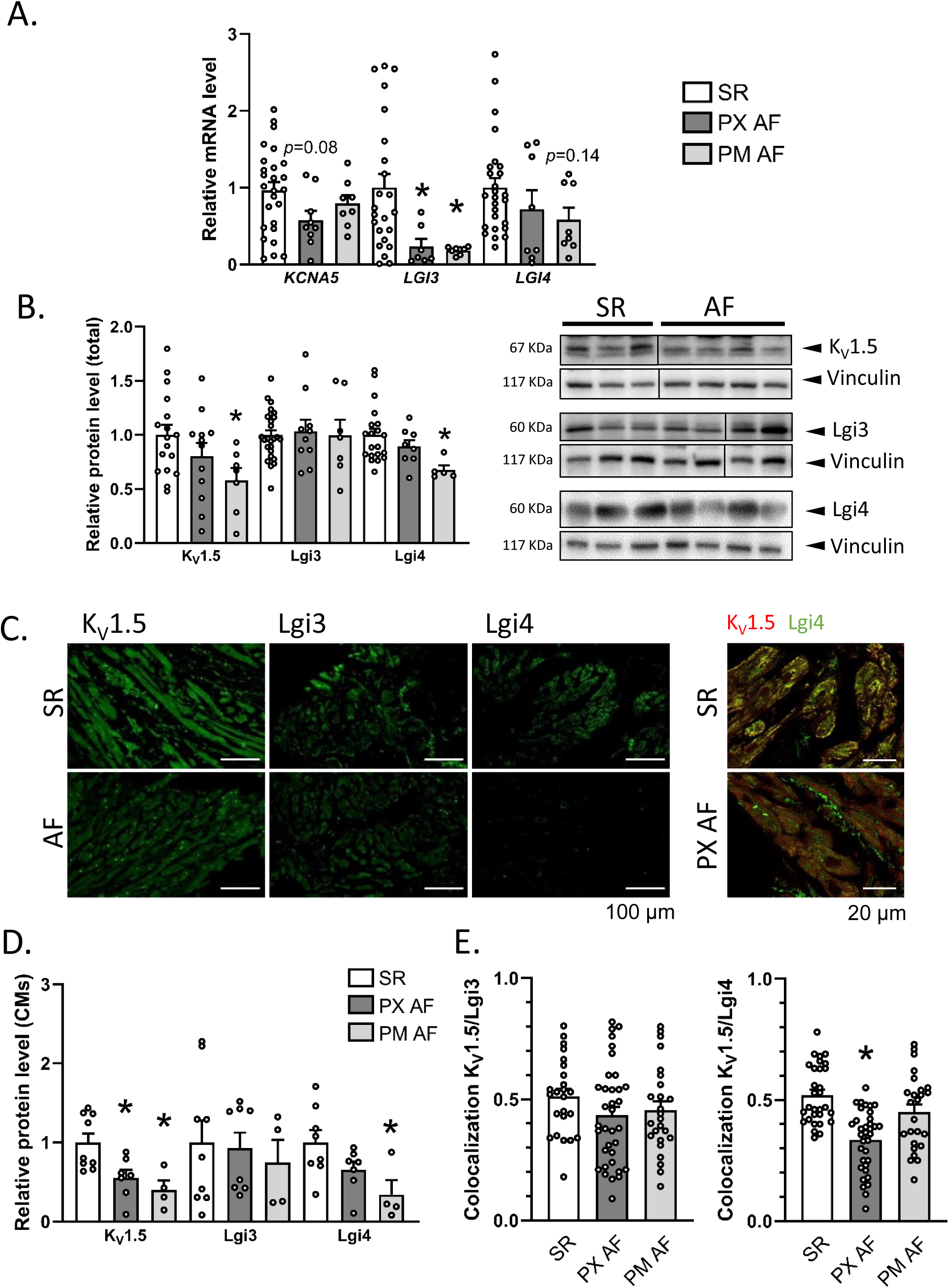
Changes in the expression of K_V_1.5 channelosome in samples from patients in atrial fibrillation (AF) compared to those in sinus rhythm (SR). **A)** Relative mRNA expression of *KCNA5*, *LGI3* and *LGI4* in right atria tissue from patients in SR and with AF determined with qPCR. AF patients are segregated on account of the progression of AF in paroxysmal (PX) and permanent (PM) AF. **B)** Relative protein expression of K_V_1.5, Lgi3 and Lgi4 in right atria tissue from patients in SR and with PX or PM AF probed in western-blot with anti-K_V_1.5, anti-Lgi3 or anti-Lgi4 antibodies and normalized by vinculin expression. Representative immunoblots using anti-K_V_1.5, anti-Lgi3, anti-Lgi4 or anti-Vinculin are shown on the right. C) *Left*-Representative confocal images of the immunodetection of K_V_1.5, Lgi3 and Lgi4 in the right atria of patients in SR (upper panels) and with AF (lower panels). Right-Representative double immunofluorescence images of K_V_1.5 and Lgi4 in SR and PX AF samples. **D)** Relative protein expression of K_V_1.5, Lgi3 and Lgi4 in atrial myocytes from patients in SR and with AF measured in individual cardiomyocytes from confocal images obtained at 40X. Each point represents the mean of 50-100 CMs from two independent IF for each patient. **E)** Colocalization of K_V_1.5 and Lgi3 (left) or Lgi4 (right) in atrial myocytes from patients in SR, PX AF or PM AF measured in individual cardiomyocytes from confocal images obtained at 63X. One-way ANOVA with Tukey test was performed when comparing more than two groups. \**p* < 0.05.

## DISCUSSION

The present work provides, to the best of our knowledge, the first study on the expression and function of Lgi proteins in the cardiovascular system. We have identified Lgi3-4 as new members of the K_V_1.5 channelosome, modulating its trafficking and biophysical properties, thereby influencing cardiac electrophysiology. These findings significantly expand our knowledge about the physiological role of these proteins. The implications are critical for understanding the macromolecular complexes that modulate *I*_Kur_, the main repolarizing current in the human atria, as well as its pathophysiological contribution to cardiovascular diseases such as AF.

Lgi1 has been associated with K_V_1 and K_V_4 channels in neurons and, more recently, Lgi3 has been linked to the trafficking of K_V_1 channels in these cells ^24,27–29,37^. These effects seem to be mainly mediated by ADAM proteins (ADAM11, 22 and 23). However, this mechanism of action cannot explain the results reported by Schulte et al., in which they demonstrated that Lgi1 abolishes the N-type inactivation induced by the K_V_β1 subunit on K_V_1.1, but not the intrinsically exerted inactivation produced by K_V_1.4 ^27^. In this study, we have demonstrated that Lgi3-4 are the only Lgi family members substantially expressing in human atrial and ventricular CMs; Lgi1-2 are virtually absent in these tissues. Moreover, Lgi3-4 interact with K_V_1.5 and K_V_4.3 channels both in cardiac tissue (in human atrium and mouse ventricle) and in heterologous systems. Such interactions seem to occur intracellularly. Our results suggest that Lgi3-4 interact with cytoplasmic K_V_β subunits, modulating their effects on K_V_1.5 channels, exerting their effects intracellularly, similarly to previous reports ^27^.

We demonstrate that one of the main effects of Lgi3-4 is the decrease of K_V_1.5 current amplitude and membrane expression when K_V_β subunits are present. Ion channel trafficking is tightly modulated to ensure proper channel function. K_V_1 channels are folded, glycosylated and assembled in the ER, and then translocated to the Golgi apparatus, where they mature before being transported to the plasmalemma ^57–59^. Such a process represents the main limiting step for trafficking/expression of most membrane proteins. K_V_β subunits enhance maturation of some K_V_1 channels ^16,60,61^ and, even though it did not seem the case for K_V_1.5 in heterologous systems, recent research demonstrated that K_V_β subunits increase *I*_Kur_ and membrane expression of K_V_1.5 in native myocytes ^15,62^. Our coimmunoprecipitation studies demonstrated that Lgi expression decreased K_V_1.5/K_V_β interaction. The fact that Lgi3-4 hinder the association between K_V_1.5 and K_V_β may suggest that the binding of Lgi proteins with K_V_1.5 and/or K_V_β occurs in the ER as an early biosynthetic event, where K_V_1.5/K_V_β interaction occurs ^60–63^. Therefore, the decrease in Kv1.5 membrane expression and current amplitude in the presence of Lgi3-4 and K_V_β subunits can be, explained, at least in part, by the decrease in K_V_1.5/K_V_β interaction. K_V_1 channels membrane expression is also modulated by internalization and intracellular retention of newly synthesized channels, but more research is needed to decipher if Lgi3-4 modulate these processes.

The reduced K_V_1.5/K_V_β interaction produced by Lgi3-4 also explains the abolishment of the N-type inactivation induced by K_V_β1 subunits on K_V_1.5, as previously reported for K_V_1.1/K_V_β1 ^27^. However, Lgi3-4 does not alter the K_V_β1-induced changes in voltage-dependence of K_V_1.5 activation and deactivation kinetics. This indicates that the interaction between K_V_1.5/K_V_β1 is sufficient to produce those effects exerted through the C-terminus of the K_V_β1, but not those induced through the K_V_β1 N-terminus. Similar patterns were observed with K_V_β1.3 N-terminus mutants and mutants in the S6 of K_V_1.5. Those variants retained their ability to cause negative shifts in K_V_1.5 activation and inactivation, even when the inactivation degree was modified, suggesting that these gating effects were mediated by interaction other than one affecting N-type inactivation ^64,65^. Perhaps, in addition to a decreased interaction between K_V_1.5 and K_V_β subunits, the interaction of Lgi proteins with K_V_1.5 and/or K_V_β subunits hinders the introduction of the K_V_β1 N-terminus into its binding site in the K_V_1.5 pore. Further, in the presence of Lgi3-4, K_V_1.5/K_V_β2.1 currents are more similar to those of K_V_1.5 alone, decreasing the C-type inactivation and shifting the midpoint of the voltage-dependence of activation towards more positive potentials. This reverts part of the effects induced by K_V_β2.1, but not those on the deactivation kinetics or the voltage dependence of inactivation. That could also be explained by a decreased interaction between K_V_1.5 and K_V_β2.1 in the presence of Lgi3-4.

Secreted Lgis, bind to ADAM23 in the nervous system. However, we have not observed any effect of ADAM23 on Lgis modulation of K_V_1.5/K_V_β1.3 currents. Hence, contrary to what occurs in neurons ^23,24,28,34,66,67^, cardiac Lgi3-4 effects are not mediated through ADAM23. This further suggests that the effects of Lgi3-4 on K_V_1.5/K_V_β are produced intracellularly. Nevertheless, ADAM23 and K_V_β proteins share some structural hallmarks, both being globular proteins with α-helical structures. This may explain the interaction of Lgi proteins with K_V_β subunits, presumably through the EPTP domain of Lgi similar to ADAM23. It also explains why ADAM23 does not induce changes in K_V_1.5/K_V_β/Lgi currents produced by intracellular Lgi protein binding to K_V_1.5 and K_V_β.

Analysis of lead-II ECG data in anesthetized mice revealed a lengthening in the QRS complex, but the rest of the parameters (P wave, PR and QTc duration) were unaltered in Lgi4 mice. In human, the QRS complex is determined by the upstroke velocity of the AP (mainly due to the *I*_Na_ amplitude); however, in mice with much shorter APs, the QRS reflects membrane depolarization (*I*_Na_), and part of the early repolarization (*I*_Kur_ and *I*_to_) of the ventricle ^53^. Therefore, the slower early repolarization observed in isolated Lgi4 CMs may explain, at least in part, the longer QRS interval on the surface ECG. Moreover, the RR variability is increased in Lgi4 with respect to control mice, thus indicating a possible alteration in the conduction system. We further investigated the effects of Lgi4 on the cardiac conduction system by challenging the heart with PES. The increase in the PR duration and concurrence of AV block after ventricular PES indicate an AP conduction alteration either through the AV node and/or the His-Purkinje system. Moreover, the increase in the SNRT following atrial PES suggests that SAN function is modified by Lgi4 expression. SAN and AVN cells express K_V_1.5 ^68–70^, and a reduction in *I*_Kur_ in SAN and AVN decreases the firing frequency of these structures ^68^, which can explain, at least in part, why Lgi4-induced *I*_Kur_ reduction may have an impact on the conduction system. Moreover, no atrial or ventricular arrythmias were reported when expressing Lgi4, which may correlate with the fact that this protein is downregulated in AF.

Early repolarization of the APs ventricular Lgi4 CMs was slower than control, and the slowing increased progressively more after changing to higher stimulation frequencies. This led to gradual AP prolongation during the application of pulse trains at frequencies higher than 4 Hz. In many CMs, this led to excessive AP prolongation with the formation of potentially arrhythmogenic EADs. Given their biophysical properties, *I*_to_ and *I*_Kur_ contribute greatly to the early repolarization of APs ^71^. The slower early repolarization process can be explained by the decreased K_V_1.5 membrane expression and *I*_Kur_ in Lgi4 CMs, which agrees with the heterologous systems data, as mice ventricular CMs express K_V_β2.1 and K_V_β1.2 subunits ^15,72,73^. K_V_β2 subunits promote K_V_1.5 plasma membrane expression in CMs, regulating the functional characteristics of *I*_Kur_.^15^ Thus, Lgi4 might interfere with K_V_1.5-K_V_β interaction by decreasing the K_V_β-induced effects on K_V_1.5 without modifying the total K_V_1.5 levels, as also observed in heterologous systems. *I*_to_ is not modified in our Lgi4 CMs experiments, but there was a tendency towards an increase. This could be due to the fact that *I*_to_ is generated by the activation of K_V_4.3/K_V_4.2 heterotetramers, with K_V_4.2 being predominant in mouse ventricular CMs. Lgi3-4 does not modify K_V_4.2 current. Another explanation may be the involvement of other K_V_4.3/K_V_4.2 modulatory subunits that are present in mouse ventricular CMs but not in CHO cells, such as KChIP2, DPP6, K_V_β2.1 and/or KCNE2 ^74,75^. The fact that Lgi3-4 increase K_V_4.3, but not K_V_4.2 current could be due to different reasons: 1) direct modulation of the ion channels; or 2) different modulatory mechanisms of these two channels, i.e., by CaMKII, which modulates K_V_4.3, but not K_V_4.2 ^76^. Anyhow, this is well beyond the scope of this study.

Isoprenaline activates β-adrenoreceptors, increasing systolic calcium to enable stronger heart contractions, and increasing K^+^ currents to limit the duration of the APs ^77^. In fact, isoprenaline increases *I*_Kur_ in the human atria ^55,78–80^. Isoprenaline does not modulate *I*_Kur_ in control, but increases it in ventricular Lgi4 CMs upon acute administration. This could be because Lgi4 CMs have less K_V_1.5 in the cell membrane, but likely retain them in intracellular compartments. As such, isoprenaline might increase channel trafficking to the membrane. In support to the idea, acute infusion of isoprenaline (1 μM) in human induced-pluripotent cell-derived CMs induced a rapid translocation of K_V_7.1 from intracellular storages to the plasma membrane ^77^. We used the same isoprenaline concentration which suggests a similar mechanism in our model. Based on that hypothesis, isoprenaline may reverse the effects of Lgi4 on K_V_1.5 current amplitude, and the increase in *I*_Kur_ can counterbalance the isoprenaline-induced increase in *I*_CaL_ ^77^. The net effect being no change in the AP shape. In contrast, isoprenaline does not increase *I*_Kur_ in control CMs, so *I*_CaL_ effects are not counteracted and AP duration is prolonged. In fact, in the presence of isoprenaline, the shape and properties of the AP are very similar in control and Lgi4 CMs, but not in its absence. The explanation would be that in the presence of isoprenaline the amplitude of *I*_Kur_ is very similar in CMs groups.

Numerous ion channels and modulatory subunits are dysregulated in AF, contributing to the electrical remodeling of the atria and thus, leading to changes in the AP shape ^3,11,81–83^. We studied the expression of members of the K_V_1.5 channelosome in right atrial appendages of patients in SR (control) and patients at two stages of AF progression (paroxysmal [PX] and permanent [PM]). Previously, it was reported that *I*_Kur_ decreases in human atrial CMs from patients in chronic AF compared with SR ^9^. Similarly, in our study, *KCNA5* mRNA levels were lower in atrial appendages from AF versus SR patients (Figure S10A) ^8,84^. However, K_V_1.5 protein expression was decreased in PM AF, and there was only a tendency towards its reduction in PX AF. The reduction in K_V_1.5 protein expression in PM AF is in agreement with previous studies ^3,8,85^, even though Brundel et al. also detected a lower K_V_1.5 in PX AF patients than SR patients ^8^. The discrepancies observed between mRNA and protein expression found in K_V_1.5 may suggest a compensatory mechanism involving post-transcriptional changes. One explanation could be that the calcium overload and structural changes that occur during AF increases the expression of proteolytic enzymes ^86,87^, both the ubiquitin proteasome-dependent pathway ^88^ and calcium-dependent cysteine proteinases (calpains) ^89^, whose activation leads to degradation of numerous other proteins ^8^, including K_V_1.5, among other ion channels ^90,91^. *LGI3* mRNA expression is decreased in AF in comparison to SR, but protein levels are not modified at any AF stage. *LGI4* mRNA is reduced in AF, similarly to what happened with Lgi4 protein expression in PM, but not in PX AF. The reduction in *LGI3* and/or *LGI4* mRNA levels may be due to an adaptive mechanism, as a decrease in Lgis would presumably lead to a normalization of *I*_Kur_. However, these changes are not translated to Lgi3 protein content, something that does happen for Lgi4. This discrepancy between mRNA and protein contents can be explained by the fact that protein levels are usually more conserved than mRNA amount, and changes in transcription are associated with translational changes with opposite effects on the final protein level, producing a buffering effect ^92^. Moreover, the positive correlation between Lgi3 and Lgi4 may suggest that they do not compensate for each other when one of them is downregulated.

Also, Lgi4-K_V_1.5 colocalization is reduced in PX AF, but not in PM AF tissue. This might explain why although K_V_1.5 protein expression is reduced in PX AF, there is no electrical remodeling at early stages of the disease ^93^ The *I*_Kur_ amplitude is expected to be similar to SR because there would be more K_V_1.5 not interacting with Lgi4, compensating for the reduced K_V_1.5 expression. However, in PM AF, both K_V_1.5 and Lgi4 are reduced, and their interaction is similar to that in SR, so *I*_Kur_ reduction would be expected.

Taking all the above information into account, Lgi4 seems to be involved in AF, exerting different modulatory roles depending on the stage of the disease. In PX AF tissue, Lgi4 interaction with K_V_1.5 is reduced, thus compensating for K_V_1.5 decrease at protein levels. On the other hand, in PM AF, both Lgi4 and K_V_1.5 are reduced, without changes in their interaction. The decrease in Lgi4 could partially compensate for the reduction in K_V_1.5 and *I*_Kur_ seen in AF. However, it would not be enough to normalize *I*_Kur_ to its SR levels. The results we have presented open up new perspectives at the pharmacological level. We demonstrate that Lgi3-4 represent new modulatory subunits of the K_V_1.5 channelosome, as well as possible new therapeutic targets towards atrial fibrillation. The expression of Lgi3-4 and their electrophysiological effects on the function of K_V_1.5 currents will allow validation of these proteins for therapeutic intervention.

## ACKNOWLEDGEMENTS

The authors would like to thank Drs. A. Felipe for providing the plasmid K_V_1.5 pCDNA3.1, Dr. Snyders for supplying K_V_1.5-GFP pBK, Dr. D. Fedida for provision K_V_1. 5-HA pcDNA3. 1, Dr. M.M. Tamkun for provide K_V_β1.3 IRES K_V_1.5 pBK, K_V_β1.2 pBK, K_V_β2.1 pBK, Dr. E. Delpón for providing K_V_4.2 GFP, Dr. F. Faivre-Sarrailh for supplying ADAM23 mCherry. We want to thank services of microscopy of the IIBm CSIC-UAM, SIDI-UAM and CNIC Viral Vectors Unit for producing the AAV9.

## FUNDING

Supported by Grants PID2019-104366RB-C21 (to C.V.), PID2019-104366RB-C22 (to M.G-R.), PID2022-137214OB-C21 (to C.V.); PID2022-137214OB-C22 (to M.G-R.) funded by MCIN/AEI/10.13039/501100011033; Grant CB/11/00222 funded by Instituto de Salud Carlos III CIBERCV (to C.V.); Grant S2022/BMD-7223 by Comunidad Autónoma de Madrid (CAM) (to C.V.); Grant FPU17/02731 (to P.G.S.); Grant BES-2017-080184 (to A.d.B.-B.); funded by Ministerio de Ciencia e Innovación. Supported also by National Heart, Lung and Blood Institute, NIH, grant number R01HL163943; La Caixa Banking Foundation project code HR18-00304 (LCF/PR/HR19/52160013); grants PI-FIS-2020 # PI20/01220 and PI-FIS-2023 # PI23/01039 from Instituto de Salud Carlos III (ISCIII) and co-funded by Fondo Europeo de Desarrollo Regional (FEDER), and by The European Union, respectively; grant PID2020-116935RB-I00 and BFU2016-75144-R funded by MCIN/AEI/10.13039/501100011033; Fundación La Marató de TV3 (736/C/2020) “amb el suport de la Fundació La Marató de TV3”; CIBERCV (CB16/11/00458; CB/11/00222 to CV); European Union’s Horizon 2020 grant agreement GA-965286; and Program S2022/BMD7229-CM ARCADIA-CM funded by Comunidad de Madrid (to JJ).

## DISCLOSURES

None

## AUTHOR CONTRIBUTION

P.G.S. and C.V. co-designed the work; P.G.S. performed most of the experiments; P.G.S, A.M. and M.R.M carried out cellular electrophysiology experiments; P.G.S, A.B.B. and M.V. performed molecular biology experiments; F.M.C. generated the mouse models and in-vivo characterization; M.J.C, E.R and L.A.D performed experiments with human samples, E.R.R, J.A.B.G and A.F.G provided human samples and clinical data; and M.G.R provided technical support, discussions and revisions; P.G.S. and C.V. co-wrote the manuscript and conceived the study; C.V. and J.J. provided supervision, funding and revisions; All authors discussed the results and commented on and approved the manuscript.

## Notes

### Competing Interest Statement

The authors have declared no competing interest.

